# Plant odour and sex pheromone are integral elements of specific mate recognition in an insect herbivore

**DOI:** 10.1101/293100

**Authors:** Felipe Borrero-Echeverry, Marie Bengtsson, Peter Witzgall

**Affiliations:** Biological Control Laboratory, Colombian Corporation for Agricultural Research (Corpoica), Mosquera, Colombia -; Chemical Ecology Unit, Department of Plant Protection Biology, Swedish University of Agricultural Sciences, Alnarp, Sweden -

**Keywords:** premating sexual communication, specific mate recognition, reproductive isolation, ecological speciation

## Abstract

Specific mate recognition strongly relies on the chemical senses in many animals, and especially in nocturnal insects. Two signal types lend to premating olfactory communication in terrestrial habitats: sex signals blend into an atmosphere of habitat odorants, where plant volatiles prevail. We show for the first time that males of the African cotton leafworm *Spodoptera littoralis* perceive female sex pheromone and volatiles of its plant host cotton as a unit, rather than as independent messages. In clean air, *S*. *littoralis* males are attracted to flawed pheromone signals, such as single synthetic pheromone components or even the pheromone of a sibling species, Oriental leafworm *S*. *litura*. Presence of host plant volatiles, however, strongly reduces the male response to deficient or heterospecific pheromone signals. That plant cues enhance discrimination of sex pheromone quality confirms the idea that specific mate recognition in noctuid moths has evolved in concert with host plant adaptation. A participation of host plant odour in sexual communication suggests that mate recognition is under natural and sexual selection. Moreover, shifts in either female host preference or sex pheromone biosynthesis give rise to new communication channels that have the potential to initiate or contribute to reproductive isolation.

## Introduction

Specific mate communication and recognition, which is shaped during adaptation to natural habitats, involves both sex signals and environmental or habitat sensory cues and is under sexual and natural selection (Paterson 1978, 1980; Endler 1992; Blows 2002; Boughman 2002; Scordato et al. 2014; Rosenthal 2017). In nature, females of phytophagous insects release sex pheromone into an atmosphere which is filled with plant volatiles.

The effect of plant volatiles on the male moth behavioural response to sex pheromone has long been investigated (Landolt & Phillips 1997; Reddy & Guerrero 2004). Perception of sex and plant volatiles typically employs discrete peripheral input channels, and two different types of insect olfactory receptors, pheromone and general odorant receptors, respectively (Krieger et al. 2004; Sakurai et al. 2004; Zhang & Löfstedt, 2015). Integration of pheromone and plant volatile stimuli has been shown to occur already at the antennal level in some species (Rouyar et al. 2015; Lebreton et al. 2017) and otherwise in the primary olfactory centre, the antennal lobe (Namiki et al. 2008; Trona et al. 2010, 2013; Chaffiol et al. 2012, 2014; Hatano et al. 2015; Ian et al. 2017).

Curiously, the behavioural consequences of blending plant volatiles with sex pheromones differ widely, between species and according to the plant chemicals investigated: plant volatiles have been shown to both synergize and antagonize the male response to sex pheromone (Dickens et al. 1990; Light et al. 1993; Yang et al. 2004; Schmidt-Büsser et al. 2009; Trona et al. 2013; Badeke et al. 2016). A tentative explanation is that plant volatiles play different ecological and behavioural roles, they signal adult feeding, mating or oviposition sites, but also plants that are unsuitable as adult or larval food. Different messages conveyed by plant volatiles would account for discrepant behavioural effects when blended with sex pheromone. This is evidenced by a response modulation according to internal physiological state in males and females of African cotton leafworm, *Spodoptera littoralis* (Lepidoptera, Noctuidae): unmated female moths are attracted to floral odorants for adult feeding, and soon after mating to cotton leaf volatiles for egg-laying (Saveer et al. 2012). Male moths respond to host leaf volatiles only prior to mating, since these probably signal rendez-vous sites (Kromann et al. 2015). Cotton leafworm moths further discriminate between volatiles of preferred and non-preferred larval food plants (Thöming et al. 2013; Proffit et al. 2015) and between volatiles from healthy and damaged cotton plants (Zakir et al. 2013a,b; Hatano et al. 2015).

That deficient plant stimuli, emitted by damaged plants or non-host plants antagonize the male response to pheromone (Hatano et al. 2015; Badeke et al. 2016; Wang et al. 2016) poses the question how deficient pheromone stimuli interact with plant volatiles. Traditionally, the effect of pheromone blend composition and quality on male attraction has been studied in the field, against a uniform plant volatile background, or in the laboratory in charcoal-filtered air, without plant volatiles. We therefore investigated the combined effect of pheromone blend quality and plant volatiles on male sexual attraction.

We show that males of *S. littoralis* best respond to a mixture of conspecific, complete sex pheromone and volatiles of the larval food plant cotton. Attraction to deficient synthetic pheromone, or to heterospecific pheromone of the sibling species *S. litura* was much reduced in the presence of cotton volatiles. Likewise, the male response to pheromone-releasing conspecific females was reduced by herbivore-damaged cotton volatiles. Our results demonstrate that mate recognition in cotton leafworm is mediated by a combination of plant volatiles and sex pheromone. This finding essentially contributes to our understanding of olfactory-mediated premating communication and has bearings on concepts of phylogenetic diversification in insect herbivores.

## Materials and methods

### Insects

An African cotton leafworm *Spodoptera littoralis* (Lepid., Noctuidea) colony was established with field-collected insects from Alexandria, Egypt and was interbred with insects from Egypt every year. Insects were raised on a semisynthetic agar-based diet (modified from Hinks & Byers 1976) under a 16L:8D photoperiod, at 24°C and 50 to 60% RH. Males and females were separated as pupae into 30 × 30 × 30 cm Plexiglas cages. Three-day-old unmated male moths were used in all bioassays.

Oriental cotton leafworm *S. litura* females were provided by Dr. Kiyoshi Nakamuta of Chiba University. *S. litura* and *S. littoralis* share pheromone components and both species show cross-attraction to their respective pheromone blends (Saveer et al., 2014).

### Plant material

Cotton seedlings, *Gossypium hirsutum* (cv. Delprim DPL 491), were grown singly in pots at 25°C and 70% RH, under daylight and an artificial light source (400 W). Cotton plants used in wind tunnel assays had 8–12 fully developed true leaves. Damaged plants were obtained by letting four, 24-h starved, fifth-instar larvae feed on the plant during 4 h prior to experiments. Larvae were removed before wind tunnel experiments. Plants used for headspace analysis and behavioural tests in previous studies were grown under the same conditions (Saveer et al. 2012; Borrero-Echeverry et al. 2015).

### Chemicals

Antennally and behaviourally active cotton volatiles (Borrero-Echeverry *et al.*, 2015) were used in experiments: β-myrcene (97% chemical purity; CAS #123-35-3; Fluka), (R)-(+)-limonene (95% chemical purity; CAS # 5989-27-5; Aldrich), (*E*)-β-ocimene (91% chemical purity; CAS #13877-91-3; Fluka), 4,8-dimethyl-1,3(*E*),7-nonatriene (DMNT) (95% chemical purity; CAS #019945-61-0; a gift from Wittko Francke, Hamburg), (*Z*)-3-hexenyl acetate (99% chemical purity; CAS #3681-71-8; Aldrich), (R)-(-)-linalool (95% chemical purity; CAS #126-90-9; Firmenich), (S)-(+)-linalool (95% chemical purity; CAS #126-90-9; Firmenich) and nonanal (90% chemical purity; CAS #124-19-6; Fluka). In addition, α-farnesene (>90% chemical purity; CAS #502-61-4; Bedoukian) and β-farnesene (>90% chemical purity; CAS #18794-84-8; Bedoukian) were included since they occur in maize headspace, another *S. littoralis* host plant (Bengtsson et al., 2006; Thöming et al., 2013).

The *S. littoralis* pheromone components, (*Z*,*E*)-9,11-tetradecadienyl acetate (Z9,E11-14Ac) (main pheromone compound), (*Z*)-9-tetradecenyl acetate (Z9-14Ac), (*Z*,*E*)-9,12-tetradecadienyl acetate (Z9,E12-14Ac), were purchased from Pherobank and (*E*,*E*)-10,12-tetradecadienyl acetate (E10,E12-14Ac) was a gift from David Hall (Greenwich, UK). Isomeric purity was >96.3% and for the dienic compounds and >99.1% for Z9-14Ac. These four components were consistently found in pheromone gland extracts (Saveer et al. 2014; El-Sayed 2017). The solvent was redistilled ethanol (Labscan).

### Wind tunnel bioassay

Wind tunnel experiments were performed in a Plexiglas wind tunnel (180 × 90 × 60 cm) following the protocol of Borrero-Echeverry et al. (2015). Briefly, males and females were kept in separate rooms to avoid pre-exposure to pheromone before experiments. One h before experiments, moths were transferred individually to 2.5 × 12.5 cm glass tubes closed with gauze. Tests were carried out between 1 and 4 h after the onset of scotophase. The wind tunnel was illuminated from above and the side (6 lux), moths were flown in a wind speed of 30 cm/s, at 24 ± 2°C air temperature and 60 ± 10% RH. Incoming and outgoing air was filtered with active charcoal. Moths for every treatment (N=50) were released individually from glass tubes at the downwind end of the tunnel. Males were scored for upwind flight over 150 cm, up to ca. 30 cm from the odour source.

Synthetic odour blends were delivered from the centre of the upwind end of the wind tunnel from a piezo-electric sprayer (El-Sayed et al. 1999; Becher et al. 2010). Samples were loaded into a 1-ml glass syringe operated by a microinjection pump (CMA Microdialysis AB, Solna, Sweden) that delivered test solutions at a constant rate of 10 µl/min through Teflon tubing into a glass capillary with a narrow, elongated tip. The capillary was attached to a piezo-ceramic disk, which produced an aerosol that was carried downwind. A glass cylinder (95 mm diameter × 100 mm height), covered by a fine metal mesh (pore size 2 mm) was placed in front of the capillary as landing platform. Live plants were placed at the upwind end of the wind tunnel. In experiments with calling, pheromone-releasing females, three calling females were placed downwind from plants in glass tubes covered at both ends with a mesh.

We tested the main pheromone compound, Z9,E11-14Ac and a 4-component synthetic pheromone blend of Z9,E11-14Ac, Z9-14Ac, E10,E12-14Ac and Z9,E12-14Ac, in a 100:30:20:4 proportion. The release rate of the main compound Z9,E11-14Ac was 100 pg/min, corresponding to the amount of pheromone emitted by calling females (Saveer et al. 2014). Males were further tested with pheromone-releasing *S. littoralis* and *S. litura* females. Single plant compounds were released at a rate of 10 ng/min, and a 4-component plant volatile blend containing nonanal, (R)-(+)-limonene, (*Z*)-3-hexenyl acetate, (*E*)-β-ocimene in a 33:12:33:23 proportion (Borrero-Echeverry et al. 2015) was also released at 10 ng/min. A 5-component plant volatile blend, mimicking herbivore damage, was formulated by adding DMNT at 10 ng/min to the 4-component blend (Hatano et al. 2015). Males were further tested with undamaged cotton plants and plants on which 5th-instar larvae of *S. littoralis* had been feeding during 4 h. Males were flown to single sources of plant volatiles and pheromones, and to combinations of both.

### Statistical analysis

Generalized linear models (GLM) with a Bernoulli binomial distribution were used to analyse behavioural data. Upwind flight was used as the target effect. Post-hoc Wald pairwise comparison tests were used to identify differences between treatments. Significance was determined at α=95%. All statistical analysis was carried out using R (R Core Team 2013).

## Results

When cotton volatiles were tested one by one, only α-farnesene elicited significant upwind flight attraction in *S. littoralis* males (z = 2.066; p = 0.039) (Table 1). All of these plant volatiles significantly reduced male attraction, when added to the main pheromone compound Z9,E11-14Ac. In stark contrast, seven of these plant volatiles did not affect male attraction when mixed with the complete, four-component synthetic sex pheromone. Only three volatiles reduced attraction when mixed with the four-component sex pheromone blend: DMNT was the strongest antagonist (z = 4.602; p < 0.001), followed by (*E*)-β-ocimene (z = 2.378; p = 0.017) and (*Z*)-3 hexenyl acetate (z = 1.799; p = 0.072) (Table 1). Larval feeding on cotton leaves strongly increases release of DMNT, which has been shown to interfere with perception of the main pheromone compound Z9,E11-14Ac (Hatano et al., 2015).

**Table 1.** Male cotton leafworm *S. littoralis* upwind flight attraction to synthetic cotton volatiles (Loughrin et al. 1995; Saveer et al. 2012; Yang et al. 2013) and sex pheromone compounds (Saveer et al. 2014; El-Sayed 2017). Single cotton volatiles were tested alone, in mixtures with the *S. littoralis* main sex pheromone compound, and with an optimized, four-component synthetic sex pheromone blend. Asterisks show significant differences between attraction to pheromone alone and pheromone blended with single cotton volatile compounds; α-farnesene was the only cotton volatile to elicit significant attraction by itself (binomial GLM and *post-hoc* Wald pairwise comparison; n = 50).

| Male upwind flight attraction [%] |  |  |  |
| --- | --- | --- | --- |
| Pheromone added | None | Main pheromone compound <sup>a</sup> | 4-Component sex pheromone blend <sup>b</sup> |
| Plant compounds <sup>c</sup> |  |  |  |
| - | - | 59 | 80 |
| α-Farnesene | 18† | 28* | 95 |
| Nonanal | 0 | 20* | 88 |
| (R)-(+)-Limonene | 8 | 23* | 78 |
| (S)-(+)-Linalool | 10 | 18* | 78 |
| β-Farnesene | 3 | 25* | 73 |
| (R)-(+)-Linalool | 0 | 0* | 65 |
| β-Myrcene | 10 | 18* | 63 |
| (Z)-3-Hexenylacetate | 0 | 10* | 55* |
| (E)-β-Ocimene | 5 | 28* | 48* |
| DMNT <sup>d</sup> | 0 | 3* | 20* |

<sup>a</sup> Z9,E11-14Ac, release rate 100 pg/min
<sup>b</sup> 100:30:20:4-blend of Z9,E11-14Ac, Z9-14Ac, E10,E12-14Ac and Z9,E12-14Ac, release rate 100 pg/min
<sup>c</sup> release rate 10 ng/min
<sup>d</sup> 4,8-dimethyl-1,3(E),7-nonatriene

This differential effect of complete vs. incomplete pheromone on male attraction, when mixed with plant compounds, was confirmed by experiments using a 4-component cotton volatile blend, instead of single cotton volatiles. Attraction to a combination of this cotton blend and the main pheromone compound was significantly reduced, compared with attraction to pheromone alone (z = 3.733; p < 0.001), while a combination of the same cotton blend with four-component pheromone did not reduce attraction (Fig. 1A). An undamaged cotton plant produced the same result: male attraction was significantly reduced to the plant in combination with the main pheromone compound (z = 2.208; p = 0.027), and not with the complete pheromone blend (Fig. 1B).

**Figure 1.**
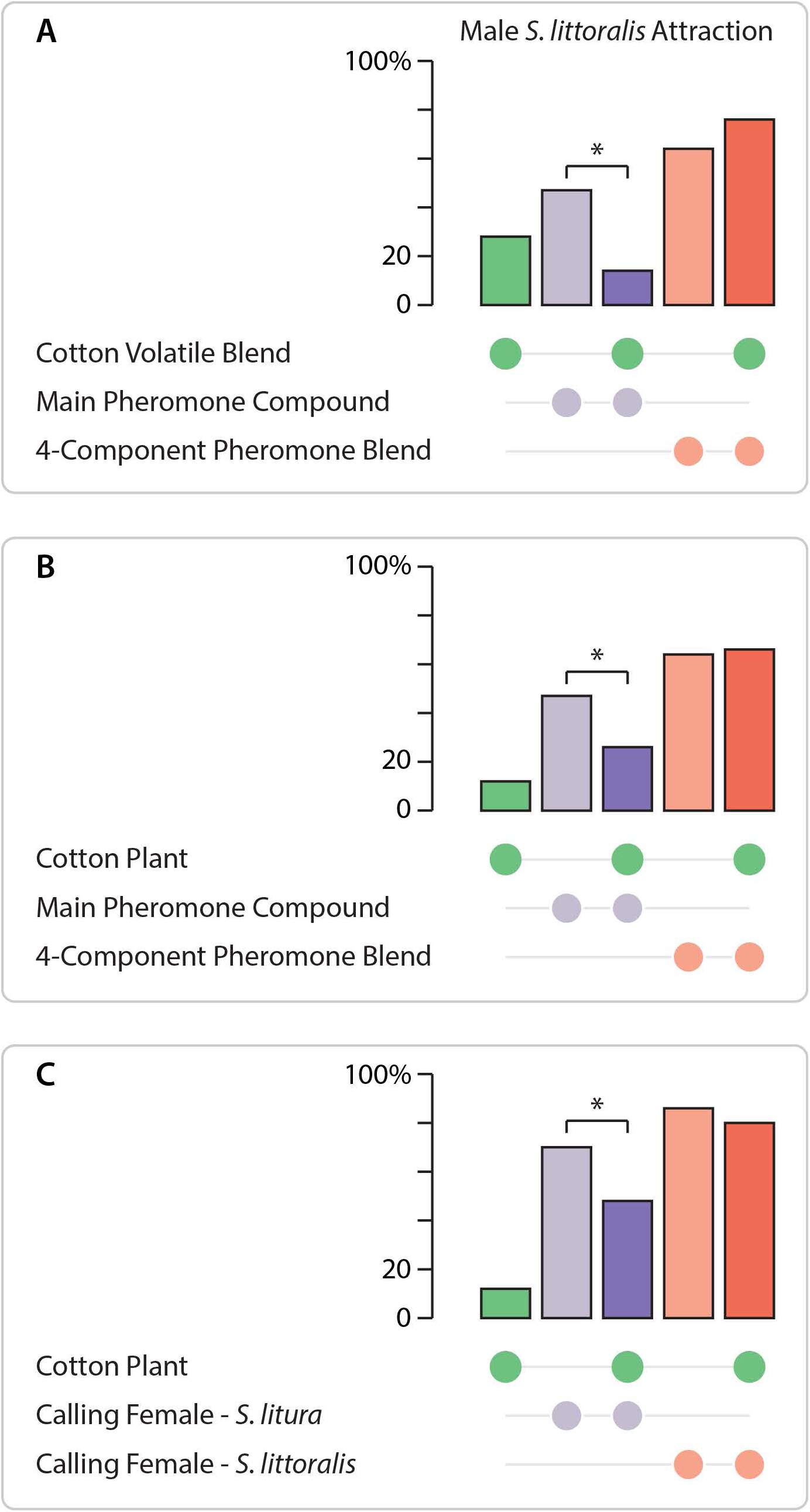
Male *S. littoralis* upwind flight attraction towards blends of *S. littoralis* females sex pheromone and cotton volatiles. Main pheromone compound alone, or optimized four-component *S. littoralis* synthetic pheromone blend (A, B), pheromone-reasing *S. littoralis* or *S. litura* females (C), in combination with a synthetic cotton volatile blend (A) or a live cotton plant (B, C). Bars with asterisks are significantly different from attraction to pheromone control (binomial GLM and *post-hoc* Wald pairwise comparisons; n = 50).

We next replaced synthetic with authentic sex pheromone, released by live calling conspecific females or by females of the sibling species *S. litura*. Both species share the main pheromone components, which explains why *S. littoralis* male attraction to pheromone-releasing females of *S. litura* and *S. littoralis* is not significantly different in clean air (Fig. 1C; Saveer et al. 2014). However, these two pheromone blends differ in composition and males are capable of discriminating conspecific from heterospecific pheromone, since they clearly prefer conspecific over heterospecific *S. litura* females in choice tests (Saveer et al. 2014). Tests with pheromone-releasing females on cotton plants confirm the results obtained with synthetic pheromone: adding cotton to *S. litura* females significantly reduced male upwind flights, compared to calling *S. litura* females alone (z = 1.992; p = 0.046) (Fig. 1C).

Lastly, we examined the effect of volatiles from cotton challenged by larval feeding on male sex pheromone attraction. We used a synthetic cotton blend and cotton plants on which *S. littoralis* larvae had been feeding. The synthetic blend mimicking damaged cotton and a cotton plant damaged by feeding larvae significantly reduced attraction to the 4-component synthetic pheromone, respectively (z = 2.208; p = 0.027 and z = 2.953; p = 0.003z = 3.899) (Fig. 2A,B). Damaged cotton plants even reduced attraction to calling *S. littoralis* females (z = 3.992; p < 0.001) (Fig. 2B).

**Figure 2.**
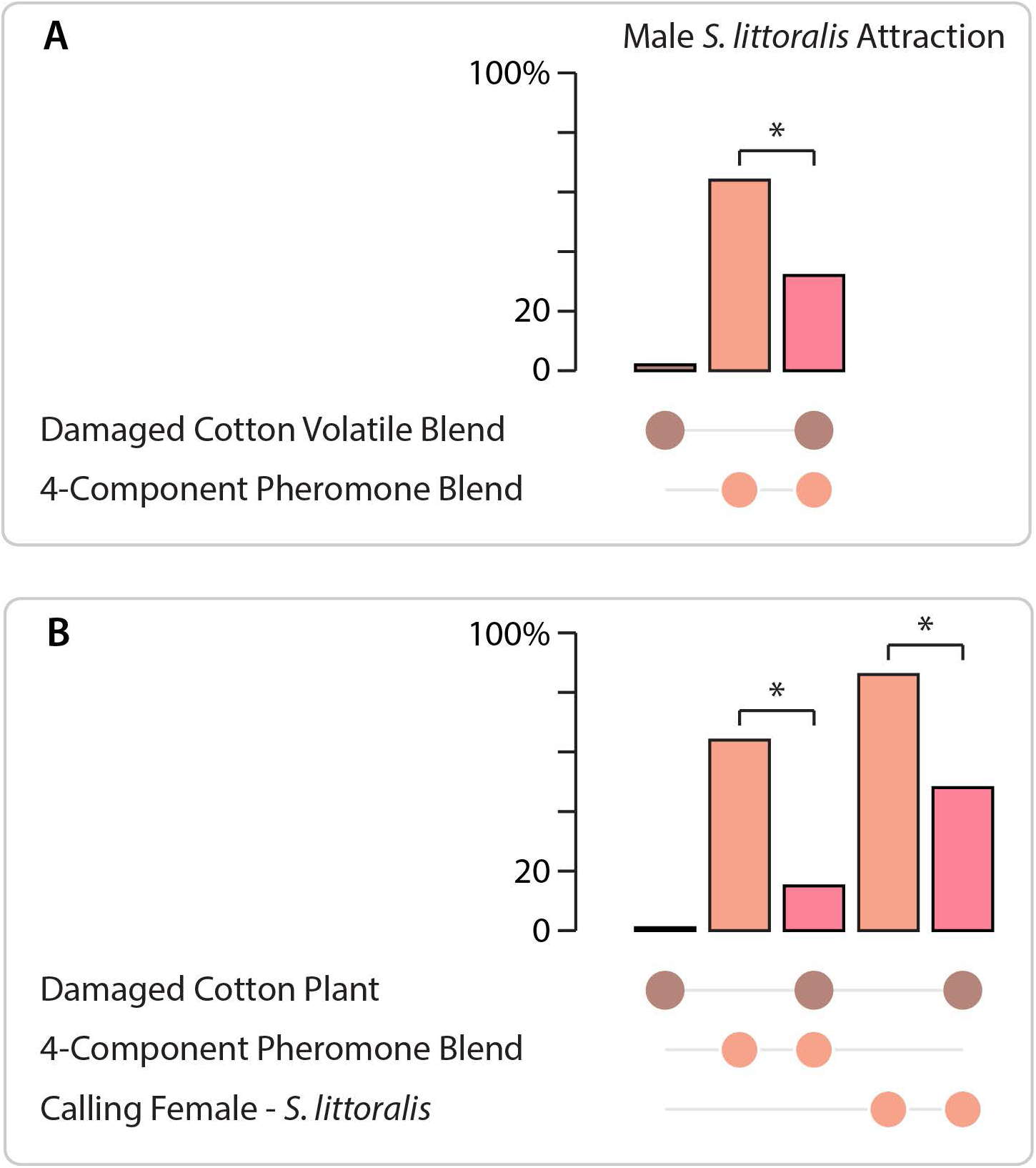
Male *S. littoralis* upwind flight attraction towards blends of *S. littoralis* pheromone and volatiles of cotton plants damaged by larval feeding. Four-component pheromone blend (A,B), or a pheromone-reasing *S. littoralis* female (B), combined with a synthetic blend of herbivore-damaged cotton volatiles (A), or a cotton plant damaged by *S. littoralis* larval feeding (B). Bars with asterisks are significantly different from attraction to pheromone control (binomial GLM and *post-hoc* Wald pairwise comparisons; n = 50).

## Discussion

That mate finding is elicited by an ensemble of sexual and environmental odorants, which mutually effect each other, provides a new perspective of premating communication in moths and has bearings for our understanding of their ecology and phylogenetic diversification.

The behavioural role of plant volatiles in male moth sexual behaviour has not been entirely resolved. It has been proposed that host plant volatiles mediate male attraction to mating sites either by themselves, before the onset of pheromone release by females, or by synergizing the response to sex pheromone (Landolt & Phillips 1997; Reddy & Guerrero 2004; Beyaert & Hilker 2014). In some species, host plant volatiles increase male attraction towards sex pheromone (Dickens et al. 1993; Light et al. 1993; Yang et al. 2004; Schmidt-Büsser et al. 2009; Varela et al. 2011; von Arx et al. 2012), whereas they produce an antagonistic effect in other species (Pregitzer et al. 2012; Jung et al. 2013; Party et al. 2013; Rouyar et al. 2015). It is conceivable that volatiles from non-host plants (Wang et al. 2016), volatiles from damaged plants, such as DMNT (Fig. 1C; Hatano et al. 2015), or floral odorants, such as β-ocimene, which signal adult food sources (Fig. 1C; Kroman et al. 2015) do not synergize pheromone attraction. It seems counter-intuitive, on the other hand, that volatiles of larval food plants would inhibit male attraction to conspecific female sex pheromone, since many moths mate on their respective host plants, where females oviposit.

Our results offer an explanation for this conundrum. Firstly, volatile signatures from damaged plants, that are less suitable for oviposition, reduce male attraction to conspecific pheromone (Table 1, Fig. 2; Hatano et al. 2015, Zakir et al. 2017). Secondly, host plant volatiles in combination with deficient or heterospecific pheromone reduce male attraction (Table 1, Fig. 1). This compares to findings in grapevine moth *Lobesia botrana*, where a blend of host plant volatiles increased male attraction to an optimized pheromone blend, but decrased attraction to a single pheromone component (Sans et al. 2016).

Cotton is a common plant host for the sibling species *S. littoralis* and *S. litura*, which are predominantly distributed in Africa and Asia, respectively. Males of the African cotton leafworm *S. littoralis* are attracted to females of both species, but prefer conspecific females in choice tests; heterospecific matings with *S.litura* females are prevented by genital morphology (Saveer et al. 2014). That presence of the plant host diminishes attraction to defective pheromone blends (Fig. 1B,C) accentuates differences between conspecific and heterospecific sex pheromones. This interaction between plant cues and sex signals is consequential, since it facilitates specific mate finding and recognition in closely related species, which frequently use pheromone blends that share compounds and partially overlap in composition (El-Sayed 2017). Sex pheromones typically consist of a blend of several compounds that have been shown to function as a unit, while the single components elicit only an incomplete behavioural response (Linn et al. 1986). Our findings demonstrate that this laboratory-derived concept must be updated to accommodate for an additional role of host plant volatiles in natural environments: it is the ensemble of social signals and environmental cues that acts as a unit to mediate mate finding and recognition.

Changes in female pheromone production and the corresponding shift in male preference are driven by sexual selection. Divergence of sex pheromone blends has been documented in populations of the same species (Cardé et al. 1978; Malausa et al. 2005; Groot et al. 2008; Velasquez-Velez et al. 2011) or in sibling species (Lofstedt & van der Pers 1985; Bengtsson et al. 2014; Saveer et al. 2014). According to the “asymmetric tracking” hypothesis, males track changes of the female pheromone composition and rather quickly develop a preference for new pheromone blends (Phelan 1992; Heckel 2010; Droney et al. 2012). Such changes are believed to enable sympatric speciation in moths through premating behavioural isolation (Smadja & Butlin 2009; M’Gonigle et al. 2012).

Host plant shifts in females, on the other hand, are driven by natural selection. If shifts in female host plant preference are disruptive, they may lead to speciation with little or no changes in pheromone composition (Lofstedt & van der Pers 1985; Witzgall et al. 1991; Drès & Mallet 2002; Bengtsson et al. 2006; Leppik & Frérot 2012). Matsubayashi *et al* (2010) suggest that changes in host plant preference may lead to premating isolation based solely on a reduced probability of encounters between populations associated with different hosts. In polyphagous species where populations have generalized diets, individuals may have preferences for a particular host plant and may be subject to selective pressures that may lead to diversification (Bolnick et al. 2003; Rueffler et al. 2006). The recent finding that host plant choice in *S. littoralis* is modified by larval experience or adult learning (Proffit et al. 2015) supports a scenario where individual preference could lead to host plant shifts and initiate divergence.

If host plant volatiles and pheromones function as a unit signal, it follows that male moth pheromone detection and mate finding is under combined sexual and natural selection. Traits combining local adaptation and mating decisions have been termed “magic traits” since they facilitate phylogenetic divergence, especially in insect herbivores (Gavrilets 2004; Smadja & Butlin 2009; Servedio et al. 2011; Thibert-Plante & Gavrilets 2013; Rebar & Rodríguez 2015). Under sympatric conditions, natural selection alone is unlikely to lead to speciation due to random mating, however, if selection acts on both habitat and mate preference simultaneously, speciation is far more likely to occur.

Our system demonstrates a mechanism where the behavioural consequences of shifts in sex pheromone biosynthesis are reinforced by host plant volatiles. New pheromone communication channels may give rise to reproductive isolation, especially in populations diverging onto new host plants. Selection acting on a combination of pheromones and host plant volatile signatures will reinforce premating barriers when a population undergoes disruptive selection (Ritchie 2007; Butlin et al. 2012; Boughman & Svanbäck 2016). This adds further support to the view that sympatric speciation has contributed to shaping the tremendous diversity of phytophagous insects (Tauber & Tauber 1989; Berlocher & Feder 2002; Drès & Mallet 2002; Forbes et al. 2017).

## Authors’ contributions

All authors contributed to designing the study and writing the manuscript. FB was responsible for insect rearing and bioassays, MB for chemicals.

### Acknowledgements

We thank Medhat Sadek and Mohamed Khallaf, Assiut, for field-collecting *S. littoralis*, and Kiyoshi Nakamuta and Saki Matsumoto, Chiba, for providing *S. litura*. This study was supported by the Linnaeus environment “Insect Chemical Ecology, Ethology and Evolution” IC-E3 (Formas, SLU), the Corporación Colombiana de Investigación Agropecuaria (Corpoica) and the Colombian Administrative Department of Science, Technology, and Innovation (Colciencias).

## Data accessibility

Data will be made available at the Dryad Digital Repository http://dx.doi.org/…

